# Fecal untargeted metabolomic and short-chain fatty acid analyses in cats with chronic kidney disease

**DOI:** 10.64898/2026.05.12.724333

**Authors:** Teresa Schmidt, Jessica M. Quimby, William H. Whitehouse, Lillian Aronson, Jan Suchodolski, Qinghong Li

## Abstract

**Background:** The gut-kidney axis plays a direct role in gastrointestinal and kidney health. Gut-derived metabolites like uremic toxins are associated with the pathophysiology of feline chronic kidney disease (CKD). The aim of the study was to identify novel fecal biomarkers and investigate the roles of gastrointestinal metabolites in feline CKD.

**Results:** Fecal samples from 41 healthy non-CKD (control) and 67 CKD cats, including 5 IRIS stage 1 (CKD1), 37 stage 2a (CKD2a), 18 stage 2b (CKD2b), and 7 stage 3 (CKD3), were subject to fecal untargeted metabolomics and targeted short-chain fatty acid (SCFA) analyses. Multiple linear regression, adjusted for sex, age, body weight and study site, identified 64 differential metabolites between control and across CKD groups (P<0.0001 and FDR<0.10). Approximately 65% of the metabolites were lipids, including polyunsaturated long-chain fatty acids, acylcarnitines, and ceramides. Random Forest algorithm selected N1-methyl-2-pyridone-5-carboxamide (2PY), a uremic toxin from nicotinamide catabolism, as the top fecal marker for classifying feline CKD. Fecal 2PY was increased in CKD1 (P = 0.03), CKD2a, CKD2b, and CKD3 (all P<0.0001) compared to the controls. Data mining revealed serum concentration of 2PY was significantly increased with severity of CKD in cats, possibly due to impaired renal excretion. Cholesterol and arachidonic acid, markers for enterocyte shedding and inflammation, were increased in CKD3 versus control (both P<0.05). In healthy non-CKD cats, evident suggested fecal lipids increased with age (P<0.0001), and were higher in females versus males (P<0.0001). While fecal indole and *p*-cresol were increased in CKD3 versus control (both P<0.05), no change was observed in indoxyl sulfate (IS) or *p*-cresol sulfate (PCS). Fecal indole-3-acetic acid (IAA) was decreased in several CKD groups compared to the controls (all P<0.05). Finally, two branched SCFAs, isobutyrate and isovalerate, were increased in CKD3 versus control (both P<0.05).

**Conclusions:** The study revealed 2PY as a novel marker and unveiled profound alterations in intestinal lipid compositions with a potential link to gut barrier integrity and inflammation in CKD.

## Background

Chronic kidney disease (CKD) affects more than 30% of cats over the age of 10 years and is reported to be the most common cause of death in cats over 5 years of age [1–4]. Typically, an initial diagnosis of CKD is made using blood and urine markers (serum creatinine, serum symmetric dimethylarginine [SDMA], urine specific gravity [USG]), followed by staging in accordance with International Renal Interest Society (IRIS) guidelines [5]. Multiple factors are implicated as risk factors that may contribute to the pathogenesis of feline CKD, but the underlying etiology remains elusive [6]. Emerging evidence highlights a significant role of the gut-kidney axis in CKD [7, 8]. Interactions between the gut and the kidney may impact the health of both organs. Recent studies revealed the occurrence of dysbiosis, an alteration of the intestinal microbiota, in a subset of CKD cats [9–11].

Intestinal microbiota exhibit an extensive metabolic function and produce metabolites, collectively known as the gut metabolome, that maintain hosts’ health or contribute to the development of diseases [12–14]. Relevant gut microbe-produced metabolites associated with kidney disease in humans and small animals include uremic toxins [9, 15–19]. Uremic toxins, including indoxyl sulfate (IS) and trimethylamine N-oxide (TMAO), are produced by intestinal bacteria during protein catabolism in the colon and accumulate in the circulation due to reduced glomerular filtration and impaired tubular secretion [8, 19–21]. Increased uremic toxin concentrations in the circulation further promote renal fibrosis, inflammation, tubular cell damage, and glomerular sclerosis [22–26]. N1-methyl-2-pyridone-5-carboxamide (2PY), a terminal nicotinamide (NAM) metabolite [27–29] and known uremic toxin, accumulates in the circulation as CKD progresses [30] and contributes to cardiovascular disease risk in humans [31]. Methods for the determination of human urinary 2PY were already developed in the late 1940s [28, 32, 33], following its isolation in urine [27]. However, reports on feline 2PY are few and far between. Other metabolic changes in CKD cats include impaired renal energetics and oxygen homeostasis [34], alterations in fecal fatty acids (e.g., short-chain fatty acids [SCFAs]) [7] and a deficiency of essential amino acids in the bloodstream [35]. Comparison of metabolite alterations between disease and healthy states can aid in a better understanding of the underlying pathophysiology of the disease, discovery of new biomarkers for diagnostics and monitoring, and identification of new potential therapeutic targets. Untargeted metabolomics are frequently used as a discovery tool to identify metabolite changes in various disease states. To date, numerous untargeted metabolomic studies using serum, plasma, and urine samples have been reported in feline CKD [34, 36–42]. Only two studies have explored untargeted fecal metabolome in feline CKD with a primary focus on dietary intervention analysis [18, 43].

The objective of this study was to identify novel disease- and stage-specific fecal metabolites that may serve as non-invasive biomarkers for feline CKD in research and clinical applications. Investigating the fecal metabolome may also provide additional insights into the gut-kidney axis and its roles in the pathogenesis of feline CKD. The findings of the current study might reveal new therapeutic targets for early intervention.

## Results

### Study population

A total of 108 cats were included in the study: 41 cats were healthy controls and 67 cats were affected by CKD, which included IRIS stage 1 (CKD1), IRIS stage 2a (CKD2a), IRIS stage 2b (CKD2b), and IRIS stage 3 (CKD3) (**Table 1**). Age and body weight were significantly different among the groups (both P<0.05). The mean age of the control group was 11.4 years compared to 13.3 years of CKD2a (P=0.04). No age difference was observed between any other groups. In addition, the average body weight of CKD3 cats was less than that of the control cats (P=0.01). No difference in body weight was observed between any other groups. Sex (**Table 1**) or study site (**Supplementary Table S1**, **Supplementary Fig. S1**) was not different among the groups.

**Table 1.**
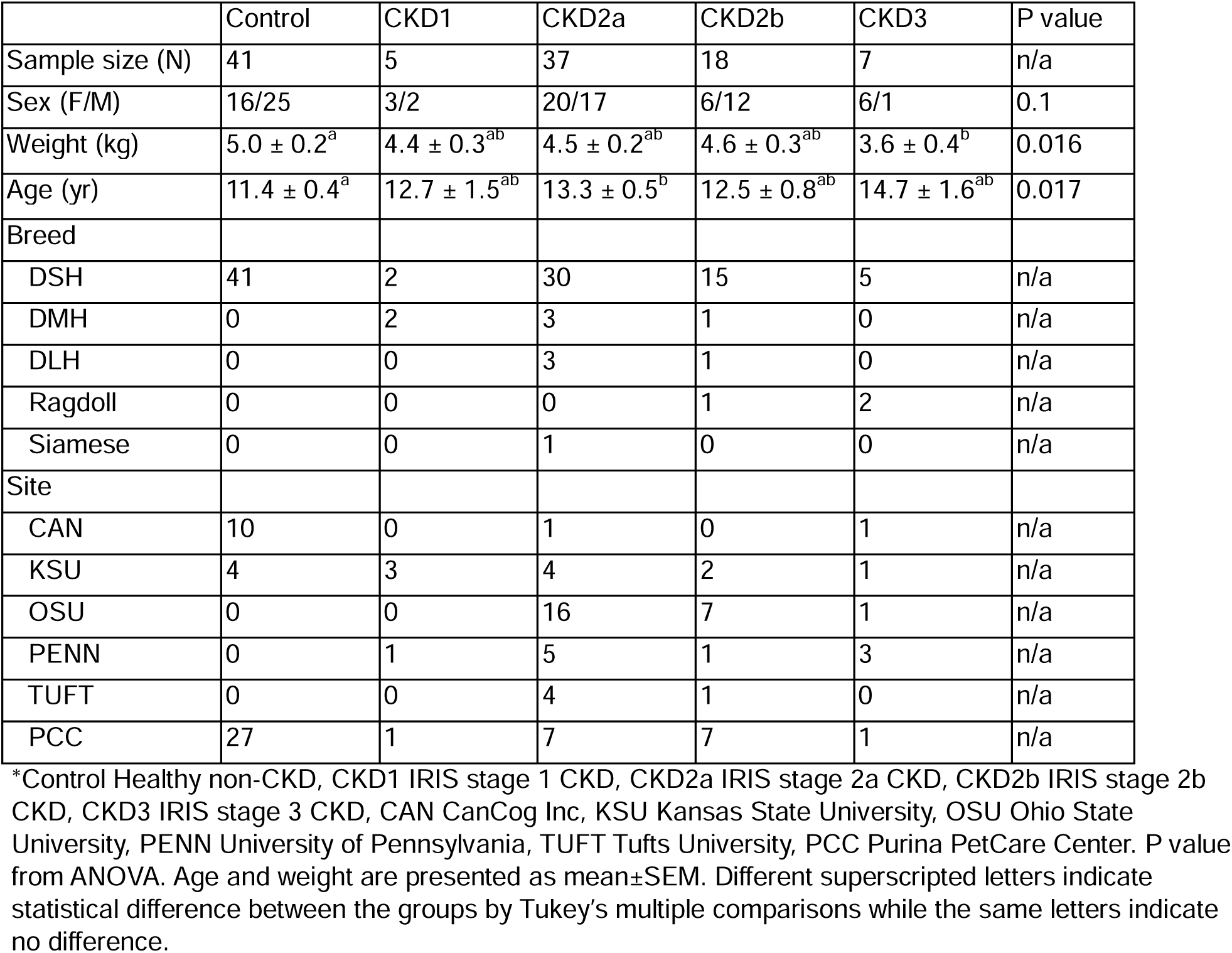
Physical characteristics of the cats and sample collection sites.

### Fecal untargeted metabolomics identified 2PY as a key fecal marker

A total of 1,369 metabolites were detected by untargeted metabolomics, 1039 (75.9%) of which had known identity (**Supplementary data file 1**). Sixty-four known metabolites were significantly different across control and different CKD groups (FDR < 0.10 and P <0.0001, **Supplementary Table S2**). The major class of metabolites was lipids (64.1%), followed by amino acids (15.6%), xenobiotics (7.8%), nucleotides (7.8%), and carbohydrates (4.7%) (**Fig. 1**). Partial Least Squares Discriminant Analysis (PLS-DA) of the 1,369 detected metabolites revealed a partial separation between healthy controls and cats affected by CKD (**Fig. 2A**). Significant overlaps were observed among the different CKD groups (**Fig. 2B**). A Random Forest machine learning algorithm was applied to the full data set to identify the best discriminative fecal metabolites separating between the control and CKD groups (**Fig. 3A**). N1-methyl-2-pyridone-5-carboxamide (2PY), a uremic toxin produced by niacin and NAM metabolism, emerged as the strongest discriminative fecal marker. Fecal 2PY concentrations were significantly increased in CKD1 (median -0.32 range [-0.8, 1.2], P=0.044), CKD2a (median 0.33 [-1.34 to 1.91], P < 0.0001), CKD2b (median 0.80 [-0.99, 1.81], P < 0.0001), and CKD3 (median 0.94 [-0.76, 2.09], P < 0.0001) compared to the control group (median -0.78 [-2.41, 1.00], P<0.0001) (**Fig. 3B**). Further, fecal 2PY showed a moderate but significant correlation with serum creatinine (Pearson r=0.51, P<0.0001, **Fig. 3C**).

**Fig. 1.**
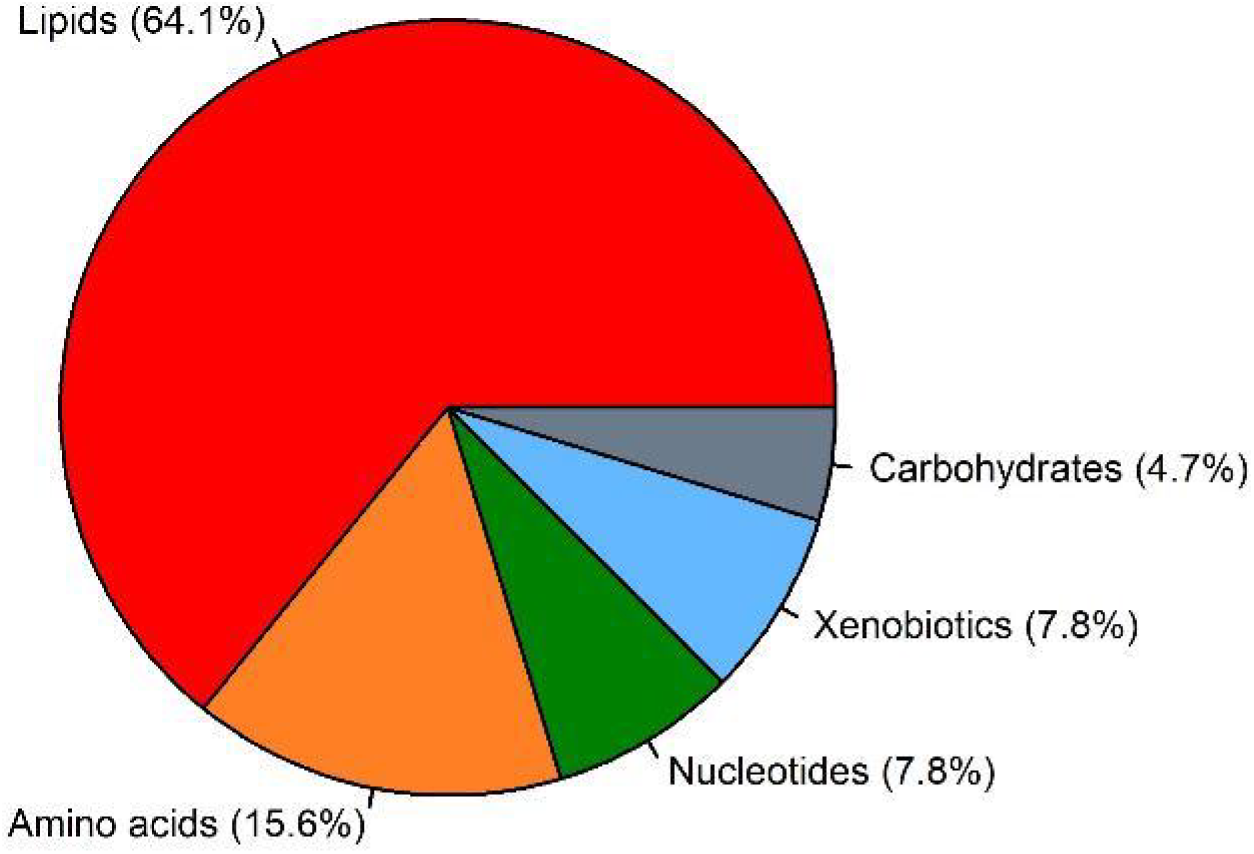
Pie chart of significant fecal metabolites. Sixty-four differential metabolites were identified. Among them, 64.1% belonged to lipids, 15.6% amino acids, 7.8% nucleotides, 7.8% xenobiotics, and 4.7% carbohydrates.

**Fig. 2.**
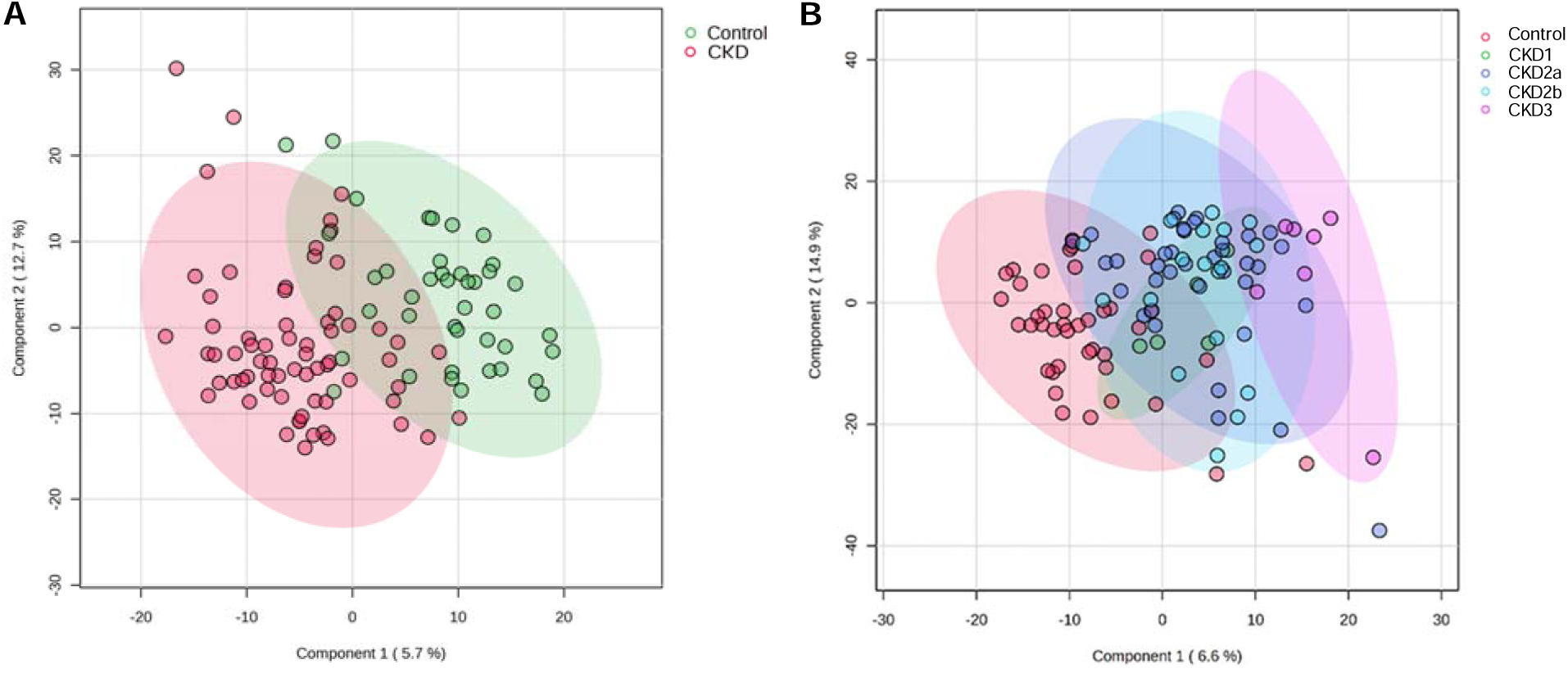
Partial Least Squares Discriminant Analysis (PLS-DA) using (A) 2 groups and (B) 5 groups. Latent variables (components) were calculated to maximize the separation among groups.

**Fig. 3.**
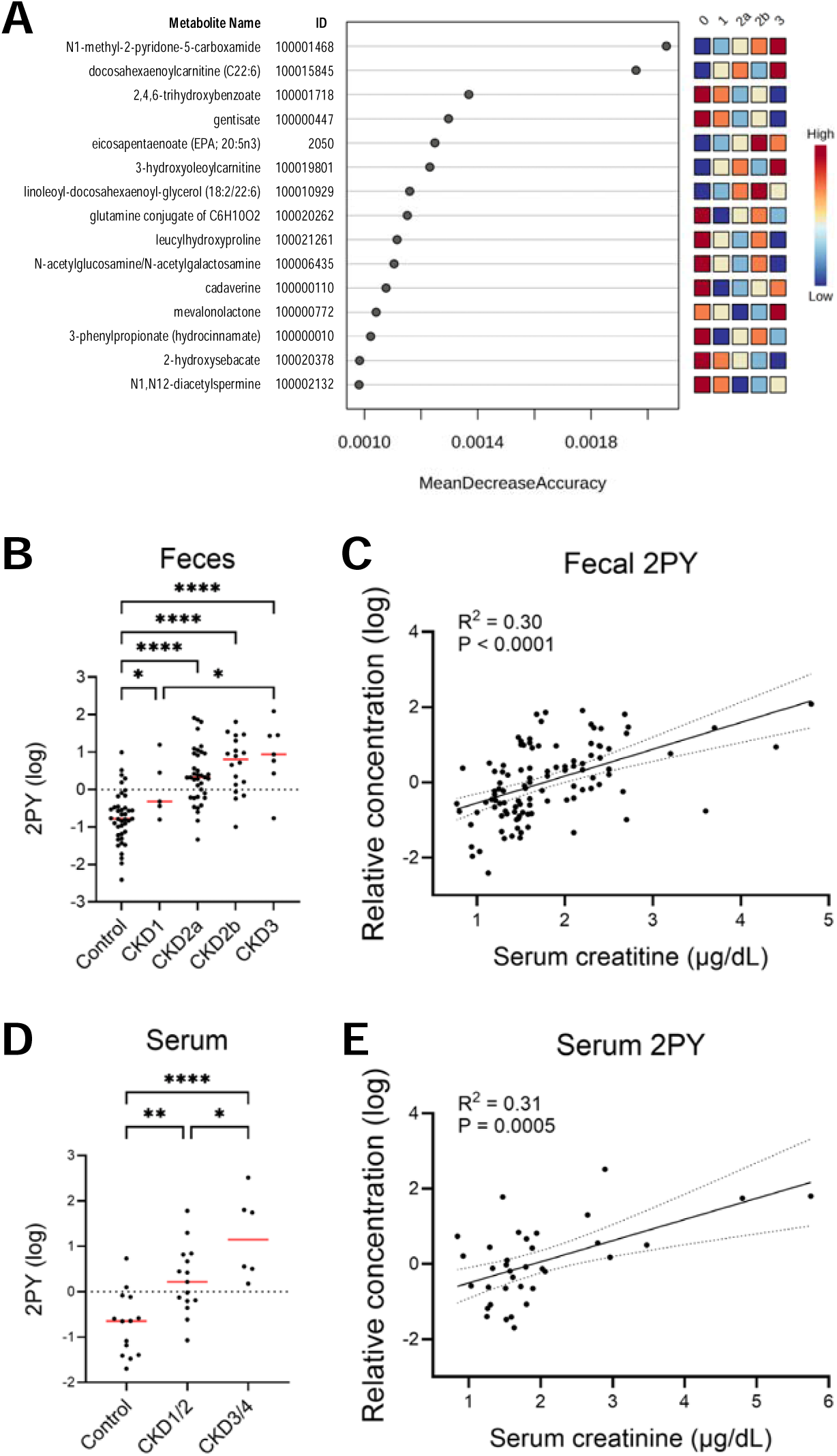
Fecal and serum N1-methyl-2-pyridone-5-carboxamide (2PY) in cats. (**A**) Top 15 features by Random Forest. Mean Decrease Accuracy (MDA) indicates individual feature’s contribution to the overall model’s accuracy. The greatest MDA was observed in 2PY. The concentration of (**B**) fecal and (**D**) serum 2PY in cats; Simple linear regression of (**C**) fecal and (**E**) serum 2PY against serum creatinine. Group means were compared using ANOVA followed by Fisher’s LSD multiple comparisons. Red lines indicate the medians. (**C**,**E**) R^2^, coefficient of determination, measures the goodness of fit for a regression model, while the P-value tests the null hypothesis that the slope is zero. The 95% confidence intervals of the estimated regression line were generated. Negative values are the results of log transformation of fractions. *P<0.05, **P<0.01, ***P<0.001, ****P<0.0001. The serum data in (**D**,**E**) were obtained from [34].

An untargeted serum metabolomic data set comparing healthy control, CKD1/2 (IRIS stages 1 and 2), and CKD3/4 (IRIS stages 3 and 4) was obtained from a previously published study [34] and the concentrations of circulating 2PY were compared among the groups. Consistent with the observation in feces, serum 2PY concentration was higher in CKD1/2 and CKD3/4 compared to healthy control (**Fig. 3D**, both P<0.01). Serum 2PY was also increased in CKD3/4 versus CKD1/2. Similarly, a moderate but significant correlation was observed between serum 2PY and creatinine (Pearson r=0.56, P<0.001, **Fig. 3E**).

### Gut microbe-produced uremic toxins and precursors

Given the central role of gut microbiota–derived metabolites such as uremic toxins in kidney disease and its progression, additional fecal metabolites were investigated. In the intestine, microbial metabolism of aromatic amino acids (tryptophan, phenylalanine, and tyrosine) produces the phenolic metabolite *p*-cresol and the indole derivatives indole and indole-3-acetic acid (IAA) [8, 20, 21]. After absorption, *p*-cresol and indole are primarily converted in the liver into uremic toxins PCS and IS, respectively, whereas IAA enters the circulation without further modification. No differences in fecal tryptophan levels were detected among groups (**Fig. 4A**). Nevertheless, IAA levels differed significantly among the groups. Cats in the CKD1, CKD2a, and CKD3 groups had lower levels of IAA compared to the control group (P=0.018, 0.012, 0.0002, respectively, **Fig. 4B**). Although fecal concentrations of *p*-cresol and indole, precursors of PCS and IS respectively, were elevated in subsets of CKD groups versus the control group **(Fig. 4C,E)**, PCS showed no differences among the groups (**Fig. 4D**). Notably, fecal IS level was below the detection limit in the majority of the samples (**Fig. 4F**).

**Fig. 4.**
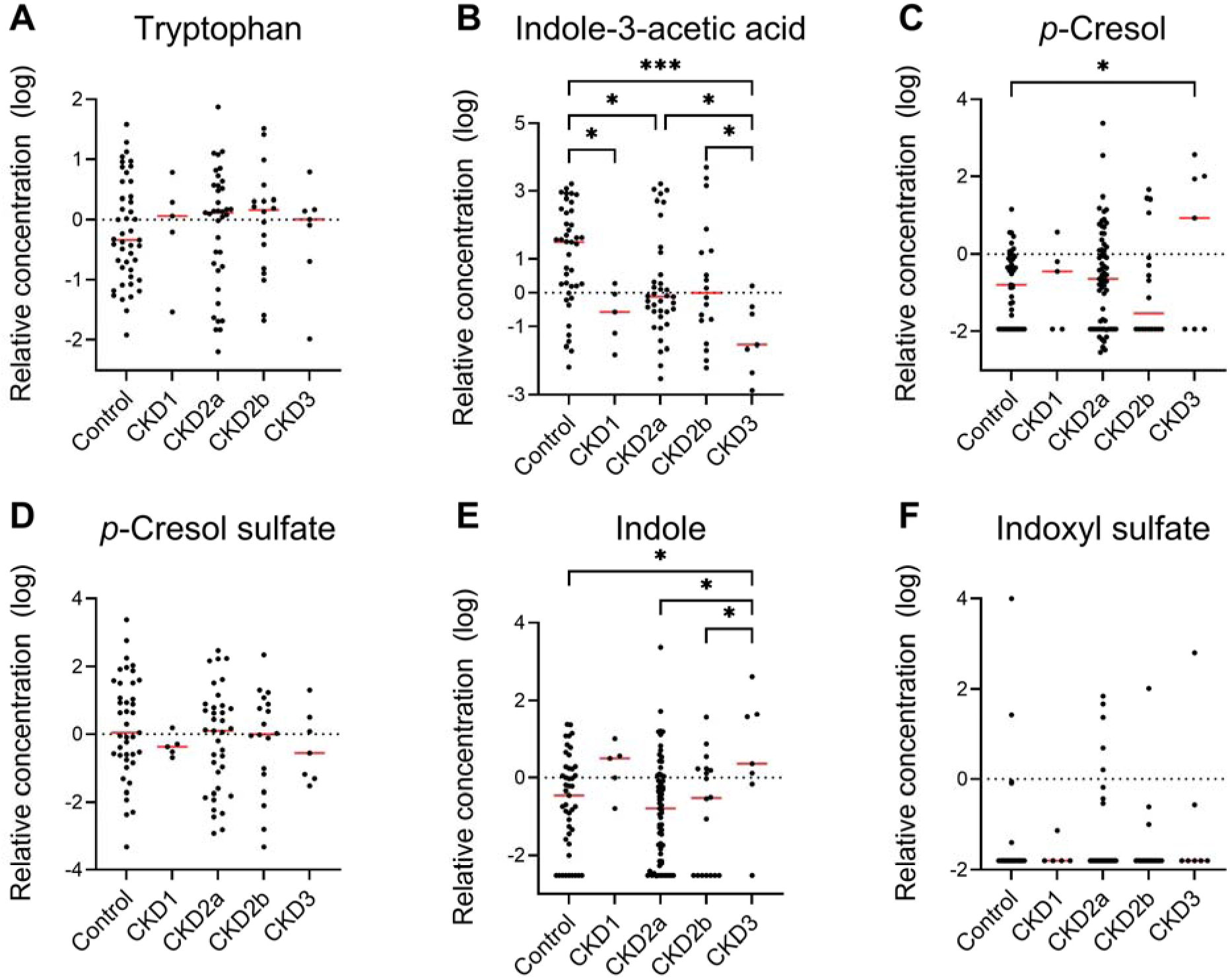
Fecal gut microbiota-derived metabolite concentrations, including uremic toxins detected by untargeted metabolomics in healthy controls and cats with CKD. (**A**) Tryptophan, (**B**) indole-3-acetic acid (IAA), (**C**) *p*-cresol, (**D**) *p*-cresol sulfate, (**E**) indole, (**F**) indoxyl sulfate concentrations across healthy controls and cats with IRIS stage 1, 2a, 2b, and 3 CKD. Red lines indicate medians. In (**F**), indoxyl sulfate was below the detection limits in the majority of the fecal samples. Significance was assessed using Fisher’s LSD multiple comparisons (* P<0.05, *** P <0.001).

### Lipids

Fecal lipids, the major metabolite group identified in the untargeted metabolomics analysis, were altered in cats with advanced CKD (**Fig. 5A**). Notably, cholesterol (CHOL), phosphatidylcholines (PC), poly- and mono-unsaturated LCFA, acylcarnitines (ACYL), and ceramides (CER) were increased in the late stages of CKD groups compared to the control group. The levels of several sphingomyelins (SPH) were increased in CKD2b, but decreased in CKD3 compared to the control. Consistently, pathway analysis comparing healthy control cats and cats with CKD identified sphingolipid metabolism as a key pathway with high statistical importance and strong metabolic pathway impact (**Supplementary Fig. S2**).

**Fig. 5.**
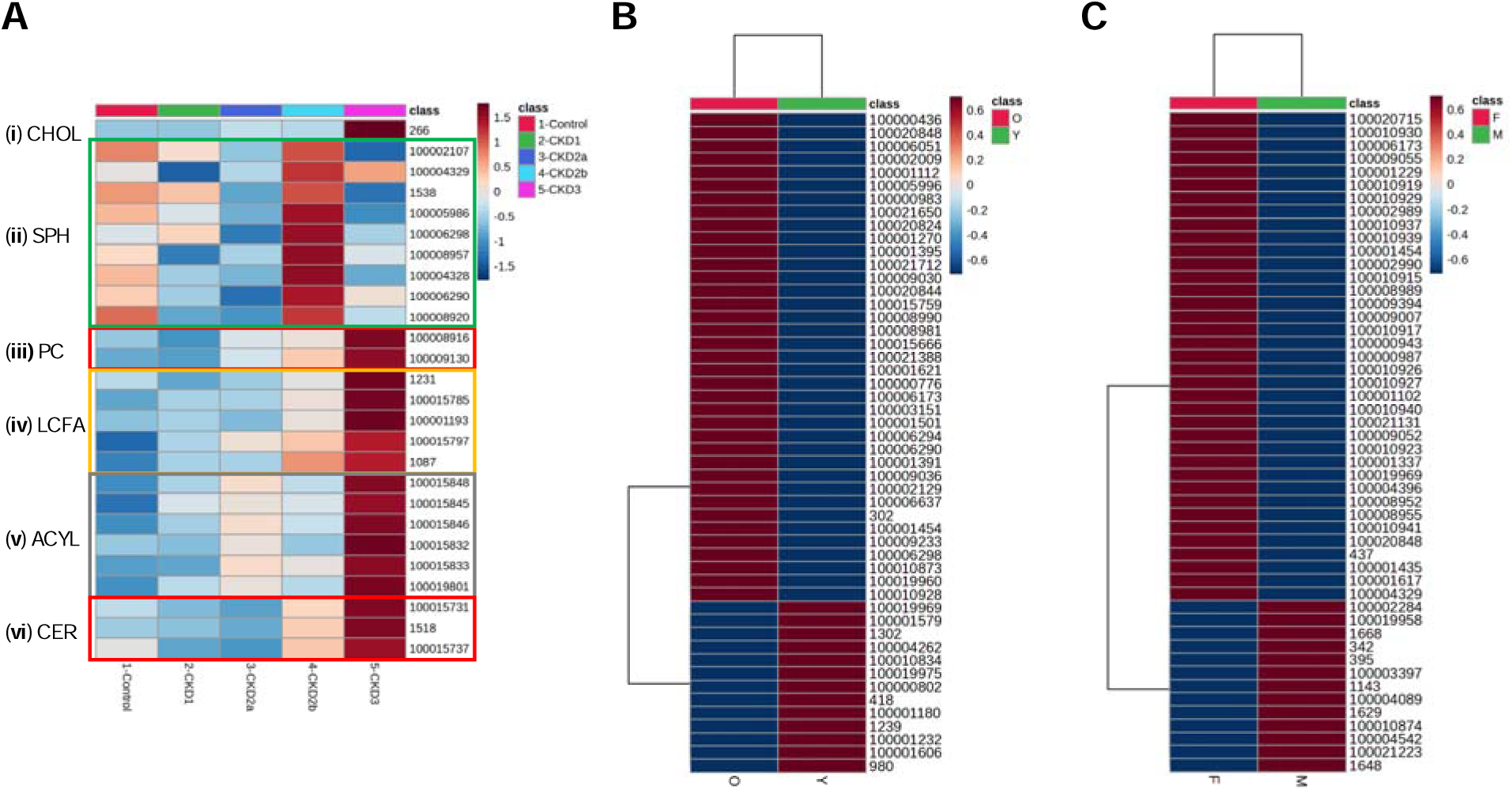
Heatmaps of fecal lipids. (**A**) The heatmap of differential lipids: (**i-vi**) cholesterol (CHOL), sphingomyelins (SPH), phosphatidylcholines (PC), poly and monounsaturated long-chain fatty acids (LCFA), acylcarnitines (ACYL), and ceramides (CER), respectively. (**B**) Top 50 lipids ranked by t-test significance between age groups: cats <11 years (Y) and cats >11 years (O). (**C**) Top 50 lipids ranked by t-test significance between males (M) and females (F). Color legend: maroon high concentration, blue low concentration. Metabolite IDs are shown to the right of eachheatmap. Group labels were displayed on the bottom of each heatmap. (**B**,**C**) Hierarchical clustering using Ward’s linkage and Euclidean distance was performed on both samples and features. (**A**) Metabolite names (from top): CHOL cholesterol, SPH [palmitoyl sphingomyelin (d18:1/16:0), sphingomyelin (d18:2/16:0, d18:1/16:1), stearoyl sphingomyelin (d18:1/18:0), sphingomyelin (d18:1/24:1, d18:2/24:0), lignoceroyl sphingomyelin (d18:1/24:0), sphingomyelin (d18:2/24:1, d18:1/24:2), sphingomyelin (d18:1/14:0, d16:1/16:0), sphingomyelin (d18:1/20:0, d16:1/22:0), sphingomyelin (d18:1/17:0, d17:1/18:0, d19:1/16:0)], PC [1-stearoyl-2-docosahexaenoyl-GPC (18:0/22:6), 1-oleoyl-2-docosahexaenoyl-GPC (18:1/22:6)], LCFA [dihomo-linoleate (20:2n6), nisinate (24:6n3), adrenate (22:4n6), heneicosapentaenoate (21:5n3), erucate (22:1n9)], ACYL [docosadienoylcarnitine (C22:2), docosahexaenoylcarnitine (C22:6), nervonoylcarnitine (C24:1), behenoylcarnitine (C22), arachidoylcarnitine (C20), 3-hydroxyoleoylcarnitine], and CER [N-palmitoyl-heptadecasphingosine (d17:1/16:0), N-palmitoyl-sphingosine (d18:1/16:0), ceramide (d18:1/17:0, d17:1/18:0)].

Further, the 41 healthy non-CKD cats were divided into two groups by age: those younger than 11 years (Y, n=18) and those aged 11 years or older (O, n=23). All 423 lipids identified in the untargeted metabolomic study were included in this analysis. Heatmaps were generated to visualize the top 50 lipids, ranked by t-test significance between the two age groups (**Fig. 5B**), and between sexes (F/M, n=16/25, all neutered) (**Fig. 5C**). The majority of these lipids (37/50=74%) has a lower concentration in the younger cats (Y) compared to the older cats (O) (**Fig. 5B**). Similarly, 37/50=74% lipids in males had a lower concentration when compared with females (**Fig. 5C**, P<0.0001). Fisher’s exact test suggested lipid level was significantly associated with age and sex (both P<0.0001).

In addition, fecal CHOL levels were significantly higher in CKD3 cats compared with any other groups (all P<0.01) (**Fig. 6A**). Similarly, fecal arachidonic acid concentrations were significantly increased in CKD3 cats compared with CKD2a and healthy controls (both P<0.05, **Fig. 6B**).

**Fig. 6.**
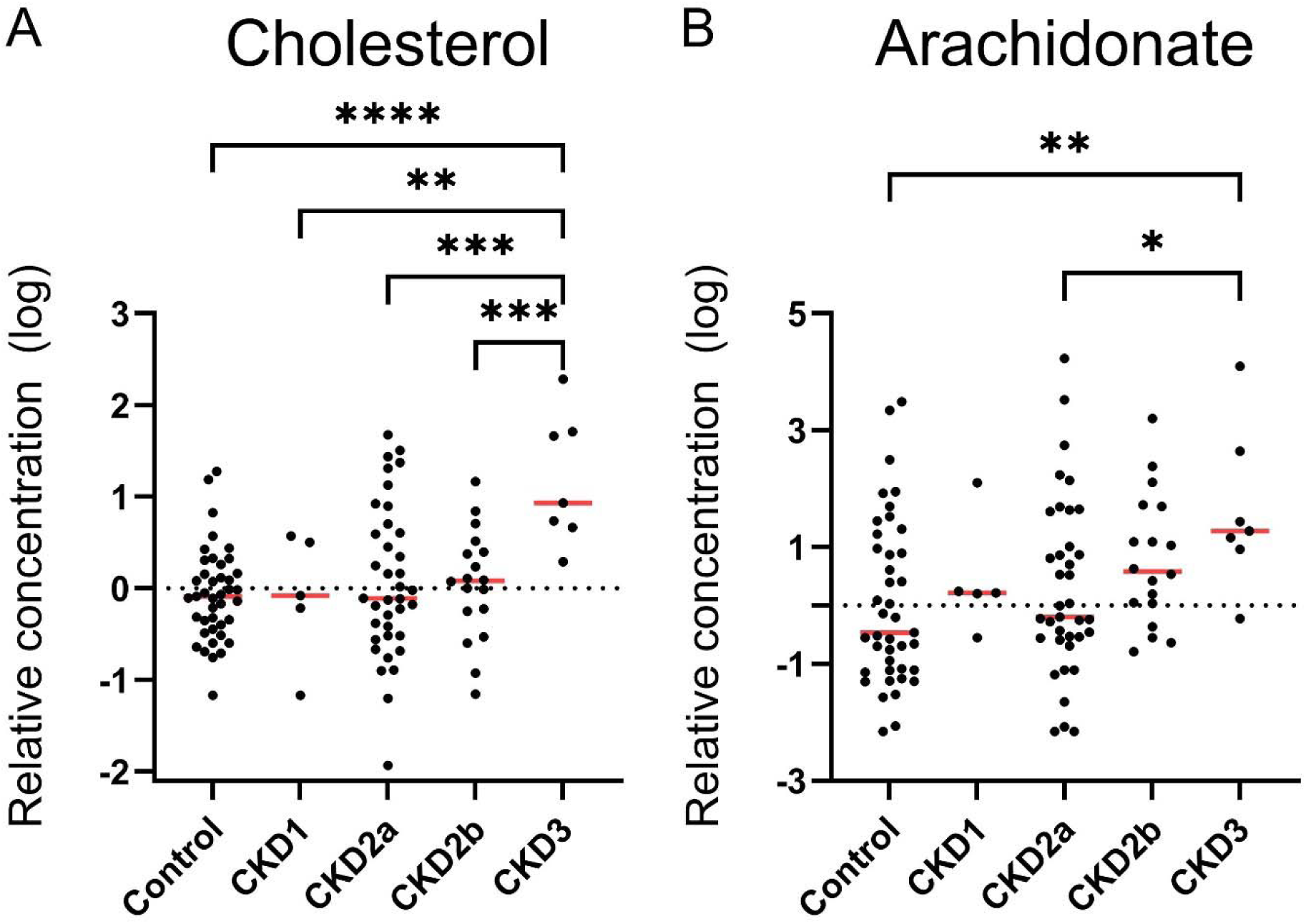
Fecal (**A**) cholesterol and (**B**) arachidonic acid. Group labels: control healthy non-CKD (n=41), CKD1 IRIS stage 1 CKD (n=5), CKD2a IRIS stage 2a CKD (n=37), CKD2b IRIS stage 2b CKD (n=18), CKD3 IRIS stage 3 CKD (n=7). Group means were compared using ANOVA followed by Fisher’s LSD multiple comparisons.

### Carbohydrates

Carbohydrate metabolism was also changed. Several fecal carbohydrates, including hexoses (glucose and fructose) and pentoses (ribose and arabinose), were decreased in CKD3 group compared to the control and CKD2b groups (**Fig. 7**). Cats in CKD3 had significantly lower fecal glucose and fructose levels compared with healthy controls (both P<0.01, **Fig. 7A,B**). Fecal fructose was also lower in CKD3 versus CKD2a (P<0.05). In addition, fecal ribose and arabinose concentrations were significantly reduced in CKD3 when compared with the control, CKD2a, and CKD2b (all P<0.05, **Fig. 7C,D**).

**Fig. 7.**
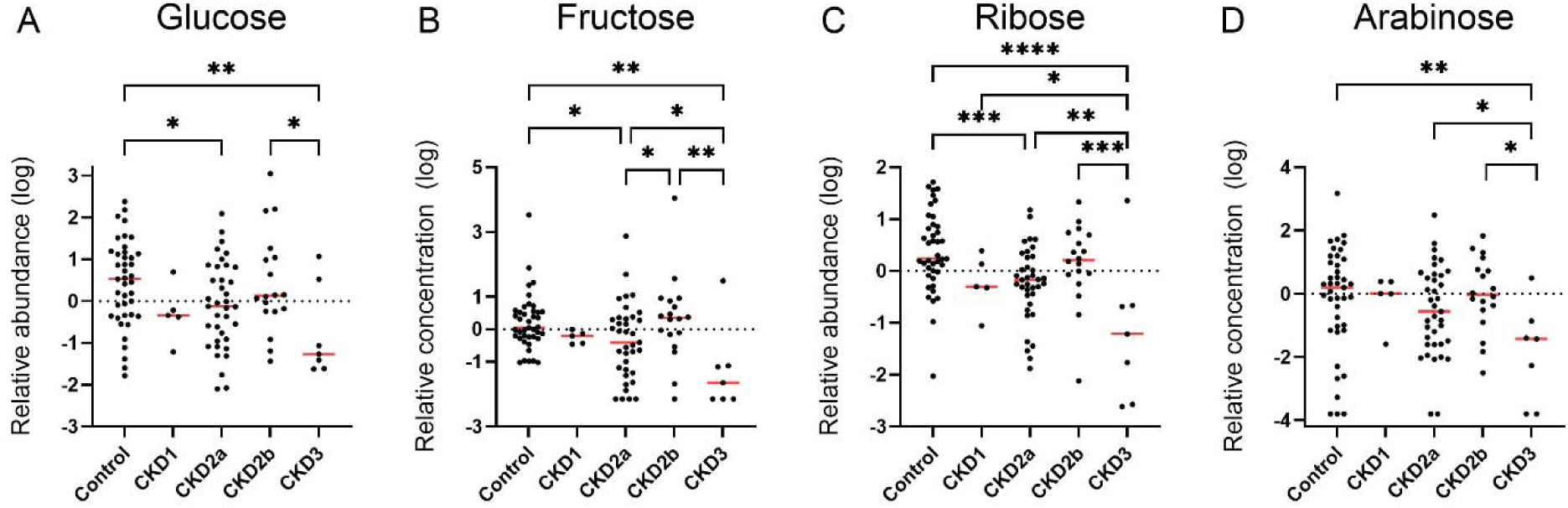
Fecal carbohydrates (**A**) glucose and (**B**) fructose, (**C**) ribose and (**D**) arabinose. Group labels: control healthy non-CKD (n=41), CKD1 IRIS stage 1 CKD (n=5), CKD2a IRIS stage 2a CKD (n=37), CKD2b IRIS stage 2b CKD (n=18), CKD3 IRIS stage 3 CKD (n=7). Group means were compared using ANOVA followed by Fisher’s LSD multiple comparisons.

### Targeted fecal short-chain fatty acid analysis

Targeted analysis revealed further alterations in fecal SCFAs in cats with CKD (**Supplementary data file 2**). Isobutyrate and isovalerate, both branched SCFAs, were increased in CKD3 compared with healthy controls (both P<0.05, **Fig. 8 A,B**). Isobutyrate was also higher in CKD3 compared to CKD2a and CKD2b (both P<0.05). No difference was observed in acetate, propionate, butyrate, or valerate (**Supplementary Table S3**).

**Fig. 8.**
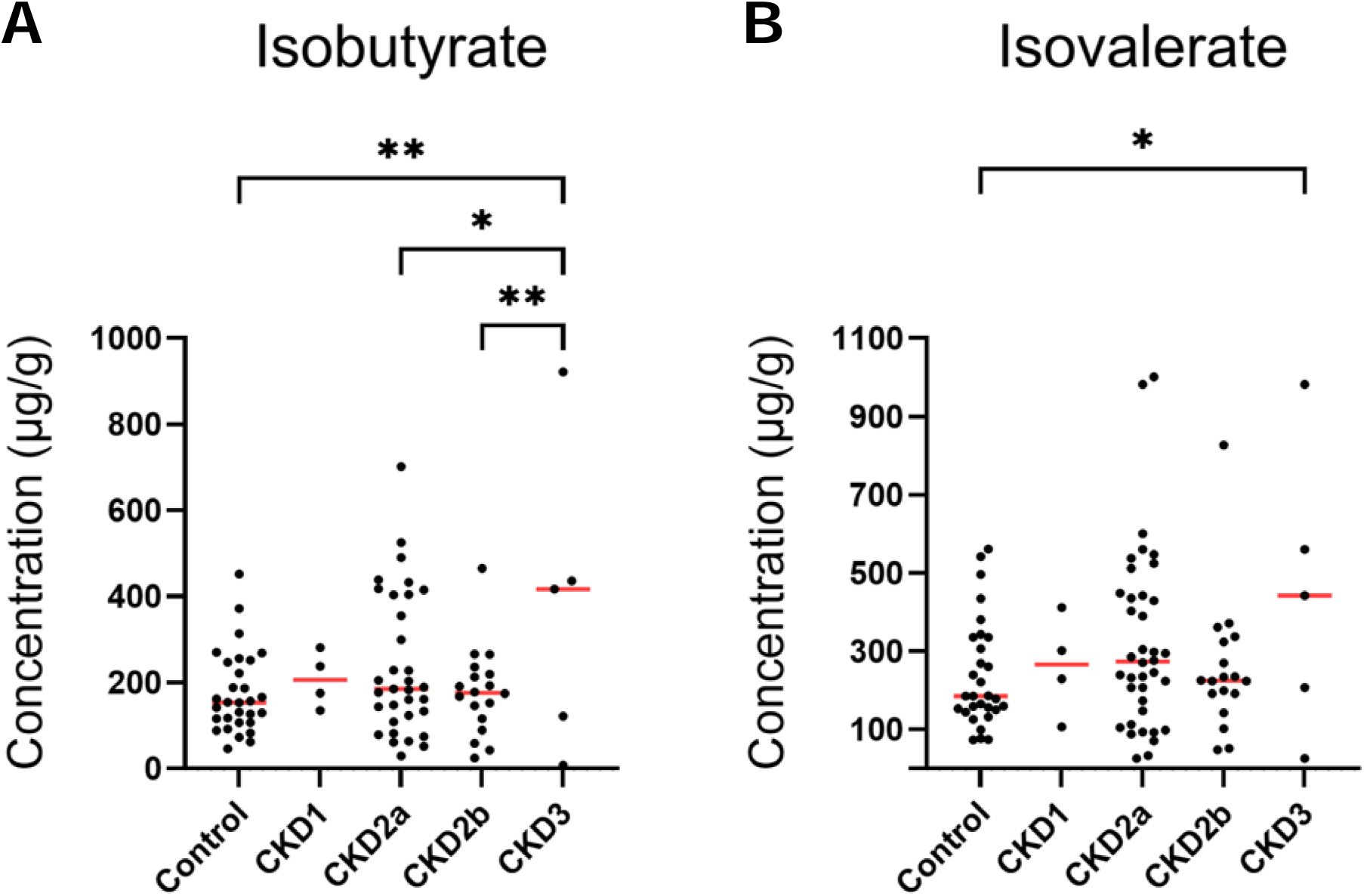
Fecal branched short-chain fatty acids (**A**) isobutyric acid and (**B**) isovaleric acid. Group labels: control healthy non-CKD (n=41), CKD1 IRIS stage 1 CKD (n=4), CKD2a IRIS stage 2a CKD (n=37), CKD2b IRIS stage 2b CKD (n=18), CKD3 IRIS stage 3 CKD (n=5). Group means were compared using ANOVA followed by Fisher’s LSD multiple comparisons.

## Discussion

In the current study, fecal samples from healthy non-CKD control cats and cats with different stages of CKD were compared using both targeted and untargeted metabolomic approaches. The aim of this study was to investigate alterations in fecal metabolites among the groups to identify novel fecal biomarkers, and potential associations between changes of intestinal metabolites and pathophysiology in feline CKD.

### N1-methyl-2-pyridone-5-carboxamide (2PY)

The strongest discriminatory fecal marker for feline CKD classification was 2PY, a uremic toxin derived from NAM catabolism [28–30]. Niacin (nicotinic acid) acts as an essential dietary precursor for nicotinamide adenine dinucleotide (NAD^+^) biosynthesis, primarily through the Preiss-Handler pathway [44]. Excess dietary niacin or NAD^+^ is metabolized to 2PY and N1-methyl-4-pyridone-3-carboxamide (4PY) via nicotinamide (NAM) catabolism [30, 45]. Although circulating 2PY and 4PY increased several-fold in both human adults and children with CKD [46, 47], a report from Yoshimura et al. demonstrated the anti-fibrotic and anti-inflammatory effects of 2PY in a mouse model of kidney fibrosis and cell culture models [48]. Recently, using untargeted plasma metabolomics and phenome-wide association analyses, Ferrell et al. revealed plasma 2PY and 4PY levels were associated with increased risk of major adverse cardiovascular events in humans [31]. Given these contradicting observations, clinical consequences of 2PY remain to be elucidated.

Literature on the concentrations and effects of 2PY or 4PY in cats is scarce, with one study reporting the daily excretion of 2PY in the urine of 2 adult male cats [49]. In this cross-sectional study, fecal 2PY concentration was increased from healthy non-CKD cats to cats with CKD and the increase was proportional with CKD progression. These observations prompted us to examine the concentration of circulating 2PY in cats with CKD. An untargeted serum metabolomic data set was obtained from a previously published multi-omics study in feline CKD [34], and serum 2PY concentrations in the CKD groups were compared to the healthy control group. Consistently, serum 2PY concentrations were several times higher in cats with the early-stage CKD (IRIS stages 1 and 2) compared to control cats, and rose even higher in cats with the late-stages (IRIS stages 3 and 4) CKD. Remarkably, both fecal and serum 2PY levels were significantly and positively associated with serum creatinine, a widely used surrogate marker for kidney function.

Accumulation of circulating 2PY is likely driven by increased NAM catabolism to remove excess niacin or NAD^+^ or by impaired renal excretion in CKD or both. The enzyme nicotinamide N-methyltransferase (NNMT) plays a key role in metabolic regulation of NAM. It catalyzes the initial methylation of NAM into N1-methylnicotinamide, which is further oxidized by aldehyde oxidase (AOX) into 2PY and 4PY. However, analyses of renal tissue gene and protein expression of AOX and NNMT showed no difference between healthy control and CKD groups in cats [34]. One cannot rule out the possibility of hyperactive NAM catabolic pathway in other tissues including the liver. Typically, 2PY, a low-molecular-weight water-soluble uremic toxin, can be easily secreted via glomerular filtration in the healthy kidneys [50]. Thus, another possibility of 2PY accumulation is due to decreased glomerular filtration rate in cats with CKD. Comparison of urine 2PY concentrations between healthy non-CKD cats and CKD cats, preferably in urine samples from timed urine collection over a 24-hour period, may shed light on the hypothesis. This study is the first to report mammalian fecal 2PY concentrations. Increases of fecal 2PY in cats with CKD raised the possibility that when renal excretion is impaired, the body may attempt to eliminate the waste products of NAM catabolism through the gut. Alternatively, the observation may also signify a “leaky” gut in CKD. Taken together, the data suggests fecal 2PY could be a novel non-invasive surrogate marker for feline CKD, which warrants future investigations.

### Fecal uremic toxins and precursors

In cats, serum concentrations of TMAO, IS, PCS, and IAA are increased in CKD [9, 19, 51]. Indoxyl sulfate has been suggested as an early marker, as higher serum abundances were detected up to 6 months before the onset of azotemia [52] and were predictive of CKD progression in cats [53]. Uremic toxin precursors indole and *p*-cresol are generated in the colon by the gut bacteria, absorbed into the portal circulation and transported to the liver, where they are converted into the final protein-bound uremic toxins IS and PCS. As these precursors are produced within the intestinal lumen by the microbiota, they are likely to be detected in the feces. In the current study, increased fecal indole and *p*-cresol were observed in IRIS stage 3 CKD, but not earlier stages, when compared with the controls. Notably, IS, the liver-sulfated indole conjugate, was below the detection limit in the majority of fecal samples. Of the three protein-bound uremic toxins, fecal PCS was decreased only in IRIS stage 3 CKD, while fecal IAA was reduced in all stages of CKD compared to the control cats. In humans, Gryp et al. reported the fecal concentration of protein-bound uremic toxins IAA, PCS, or IS was not different among healthy people and patients with different stages of CKD in an *ex vivo* fecal culture study [54]. More studies are needed in order to determine the possibility, if any, of increased uremic toxin influx from the blood into the gut due to the elevated circulating uremic toxin concentration in cats.

### Fecal lipids

Approximately 64% of altered fecal metabolites were lipids, including LCFAs, CHOL, SPH, and CERs. Fecal arachidonic acid and cholesterol were increased in cats with late-stage CKD. As both compounds are major constituents of cellular membranes, their increased fecal abundance may suggest gastrointestinal mucosal injury with enhanced enterocyte shedding associated with CKD. In healthy non-CKD cats, exploration of lipid distribution suggested that fecal lipids increased with age and were higher in females versus males. It is possible that older healthy cats also had increased enterocyte shedding. Alternatively, while a direct pathway for fat excretion from kidney-to-gut is not established in cats, a shift in lipid metabolism with age may offer a plausible explanation. Advancing age is one of the key risk factors for kidney disease [2, 55]. In humans, kidney function declines with normal aging [55, 56]. There is also evidence of renal aging and histopathologic lesions in cats without kidney disease [57, 58]. Circulating uremic toxins increase in cats older than 12 years compared with younger ones independent of CKD [19]. Intracellular lipid accumulation, particularly in proximal tubular epithelial cells, is associated with aging in normal cats and is significantly more common than other species [59–61]. Ectopic lipid deposition, within the interstitial space, increases with age and appears to be associated with inflammation and tubular loss in cats without CKD [57, 59]. Interstitial lipid may be an important driver of the development of tubular interstitial inflammation and fibrosis in the feline kidney [57]. Characterization of feline renal lipid reveals novel lipids found only in domestic cats, raising questions regarding species-specific lipid metabolism and susceptibility to disease [61]. Lastly, it is worth noting that 420 fecal lipids identified in this study, while informative, constitute only a small portion of the total fecal lipidome in cats. Thus, the findings from the study should be viewed as preliminary and warrant future investigations.

### Fecal carbohydrates

In contrast to lipids, several carbohydrates were decreased in fecal samples of cats with CKD. These findings can be interpreted in different ways. Firstly, nutrition-related factors could be one possible explanation. Cats with CKD commonly show reduced appetite, intermittent anorexia, and reduced food intake, which could lower the amount of dietary carbohydrates. In addition, renal diets differ in macronutrient composition, which could further influence fecal carbohydrate patterns independent of disease mechanisms. Secondly, decreased fecal carbohydrates may also reflect altered intestinal absorption and transport, which would be compatible with CKD-associated gastrointestinal involvement. Altered fecal profiles of transporter-dependent carbohydrates have been linked to chronic enteropathy in a subset of dogs and cats with dysbiosis, and a reduced density of carbohydrate transporters in intestinal crypts has been reported in dogs with chronic enteropathy [62, 63]. Although these changes in chronic enteropathy are typically associated with malabsorption and increased fecal carbohydrates, they illustrate how intestinal dysfunction can affect fecal carbohydrate signatures. Thirdly, lower fecal carbohydrate levels could also indicate a shift in the intestinal microbiome, reflecting increased microbial utilization of available carbohydrates. This would align with CKD-associated dysbiosis that has been observed in a subset of cats. Overall, reduced intake, altered absorption, and increased microbial utilization may overlap, and the present dataset cannot resolve their relative contributions. Further studies are warranted to clarify the drivers of decreased fecal carbohydrates in feline CKD.

### Targeted fecal SCFAs analysis

In the targeted fecal SCFA analysis, isobutyrate and isovalerate, both branched SCFAs (BCFAs), were increased in a subset of cats with stage 2 or stage 3 CKD. These BCFAs, which constitute a small fraction of total SCFAs, are produced via microbial fermentation of the branched-chain amino acids valine, leucine, and isoleucine (e.g., by *Bacteroides spp.* and *Clostridium spp.*) [64, 65]. Fecal BCFA production depends on protein availability in the colon, as only protein that escapes digestion and absorption in the small intestine becomes available for microbial fermentation [7, 66–68]. BCFAs have been associated with potentially negative effects on intestinal health, including promotion of intestinal inflammation and reduced gut motility [67–69]. In contrast, straight-chain SCFAs (acetate, propionate, and butyrate) are generally linked to beneficial effects, including anti-inflammatory properties, support of gut-barrier function, and regulation of luminal pH [70]. In humans with CKD, declining glomerular filtration rate has been associated with decreased fecal straight-chain SCFAs, correlating with reduced abundance of SCFA-producing bacteria [71]. To the authors’ knowledge, fecal BCFAs have not been specifically studied in people with CKD. However, limited feline data on BCFAs are available and are consistent with our findings. Summers et al. reported higher fecal isovalerate concentrations in cats with late-stage CKD (IRIS stages 3–4), although isobutyrate was not significantly different [7]. Notably, fecal total BCFAs, isobutyrate, and isovalerate were significantly increased in CKD cats with muscle atrophy compared with CKD cats with normal muscle mass [7].

Taken together, increased fecal BCFAs in a subset of late-stage feline CKD may reflect a shift toward proteolytic fermentation, potentially influenced by diet composition, nutrient digestion/absorption, dysbiosis, and/or CKD-associated muscle loss. Because diet, muscle condition, and microbiome profiles were not assessed in the present study, the drivers and clinical significance of increased fecal BCFAs in feline CKD warrant further investigation.

### Limitations

This study has some limitations that should be considered when interpreting the results. The study population differed between groups. Cats with CKD and healthy non-CKD controls showed a statistically significant age and weight difference. Although the control group consisted of cats aged 6 years or older, age-related effects on the fecal metabolome cannot be ruled out. In addition, sample sizes were small for IRIS stage 1 (CKD1) and stage 3 (CKD3) groups. Fecal metabolite profile may be influenced by multiple factors including diet. For example, specially formulated therapeutic diets are prescribed to feline renal patients to manage symptoms and support kidney function. It is common that cats with CKD experience decreased appetite, nausea, and weight loss. To maintain body weight and hydration, different food types, including wet food, are offered to encourage consumption and boost hydration. Medications, including those for CKD management, can also influence global metabolome change. Although cats with major systemic diseases were excluded, it was not possible to rule out cats with subclinical conditions, such as subclinical cancer or heart disease. Untargeted methods are designed for discovery purposes and the effect sizes are not known. Targeted validation is warranted for precise, reproducible measurements and to mitigate potential false positives. Paired fecal and serum metabolomics, and microbiome profiling as well as clinical markers of gastrointestinal injury are crucial in advancing our understanding of the gut-kidney axis in feline CKD. The integrated multi-omics approach will shed light on how gut dysbiosis links to metabolic abnormalities and CKD pathophysiology, offering new insights for therapeutic interventions.

## Conclusion

Overall, the findings provide new insights into the gut–kidney axis and suggest that feline CKD is associated with fecal metabolomic alterations, particularly in advanced CKD stages. Fecal 2PY emerged as the strongest discriminatory metabolite and increased progressively with CKD, suggesting that 2PY may represent a novel non-invasive biomarker for feline CKD. In addition, increased fecal lipids, such as arachidonic acid and cholesterol, suggest a potential role of gut-kidney axis in feline CKD. Changes in microbiota-related metabolites further support this potential role of microbial dysmetabolism in feline CKD. Increases in fecal indole, *p*-cresol, and BCFAs (isobutyrate and isovalerate) in late-stage CKD cats are consistent with the observation that CKD is associated with altered microbial metabolism and a shift toward proteolytic fermentation. Future studies including paired fecal and serum metabolomics, and microbiome profiling as well as clinical markers of gastrointestinal injury are warranted. Such data may help design strategies for dietary and pharmaceutical interventions to modulate gut microbial metabolism and uremic toxin burden in feline CKD.

## Methods

### Study population and design

A prospective multicenter cross-sectional study was conducted in cats affected by CKD and healthy colony cats serving as the control group. Cats were enrolled between 2019 and 2023 at six different study sites (**Table 1**). The study was approved by the Institutional Animal Care and Use Committee (IACUC) of each participating site prior to initiation. Informed owner consent was obtained for client-owned cats.

In cats diagnosed with CKD, clinical history was evaluated, physical examination, blood analysis (hematology, serum chemistry, SDMA, thyroxine [T4]), indirect systolic blood pressure (sBP), urinalysis (BP), including USG and urine protein to creatinine ratio (UPC) were performed. Kidney ultrasound was also available in a subset of cats. The cats in the CKD group was then staged according to the IRIS guidelines. A CKD cat was assigned IRIS stage 1 (CKD1) if its serum creatinine concentration was <1.6 mg/dL, otherwise IRIS stages 2-4 as appropriate. In the current study, cats in IRIS stage 2 were further sub-classified into IRIS 2a (CKD2a, creatinine 1.6-2.3 mg/dL, both inclusive) and IRIS 2b (CKD2b, creatinine 2.3-2.8, including 2.8).

Enrollment criteria for CKD included a clinical history and physical exam supporting the diagnosis of CKD, a USG <1.035, and abnormal serum creatinine or abnormal SDMA levels on at least two different assessments. Cats diagnosed with CKD with regulated hyperthyroidism and systemic hypertension were allowed to enroll. Cats with a major systemic disease such as cancer, heart failure, and diabetes mellitus were excluded.

The healthy control group consisted of healthy cats meeting the following inclusion criteria: (1) USG ≥1.035, (2) serum SDMA <18 μg/dL, (3) serum creatinine <1.6 mg/dL, (4) sBP <160 mmHg, and (5) no clinical evidence of kidney disease, urinary tract infection or any other major systemic diseases such as cancer, heart disease, or diabetes mellitus at the time of examination.

### Fecal sample collection, storage, and shipment

Fecal samples from colony cats were collected within 30 minutes of defecation and immediately stored at -80°C until use. For privately owned cats, a piece of fresh fecal matter was collected from a litter box in a sealed bag and refrigerated as soon as possible. The sample was then delivered to the veterinary clinic within 24 hours and stored in -80°C until use.

### Untargeted fecal metabolomics

Untargeted metabolomic analysis was performed in a commercial laboratory (Metabolon, Inc. Morrisville, NC). Fecal samples were prepared using the automated MicroLab STAR® system from Hamilton Company. Several recovery standards were added prior to the first step in the extraction process for QC purposes. To remove protein, dissociate small molecules bound to protein or trapped in the precipitated protein matrix, and to recover chemically diverse metabolites, proteins were precipitated with methanol under vigorous shaking for 2 min (Glen Mills GenoGrinder 2000) followed by centrifugation. The resulting extract was divided into multiple fractions: two for analysis by two separate reverse phase (RP)/UPLC-MS/MS methods with positive ion mode electrospray ionization (ESI), one for analysis by RP/UPLC-MS/MS with negative ion mode ESI, one for analysis by HILIC/UPLC-MS/MS with negative ion mode ESI, while the remaining fractions were reserved for backup. Samples were placed briefly on a TurboVap® (Zymark) to remove the organic solvent. The sample extracts were stored overnight under nitrogen before preparation for analysis.

The raw data were generated based on the area under the curve formula using ion counts that provide relative quantification. For each metabolite, the raw values in the experimental samples are divided by the median of those samples in each instrument batch, giving each batch and thus the metabolite a median of one. For studies containing <144 total samples, typically no batch normalization is required and the batch-normalized data simply reflect median-scaled raw data. Missing metabolite values can typically be attributed to the signal falling below the limit of detection. For each metabolite, the minimum value across all batches in the median scaled data is imputed for the missing values. The batch-normalized and imputed data was transformed using the natural log.

### Targeted fecal short-chain fatty acid analysis

Feces samples, including 30 healthy control, 4 CKD1, 34 CKD2a, 18 CKD2b, and 5 CKD3, were analyzed for six short chain fatty acids acetic acid (C2), propionic acid (C3), isobutyric acid (C4), butyric acid (C4), isovaleric acid (C5), and valeric acid (C5) by LC-MS/MS also by Metabolon, Inc. (Morrisville, NC). Samples were spiked with stable labelled internal standards, homogenized, and subjected to protein precipitation with an organic solvent. After centrifugation, an aliquot of the supernatant was derivatized and injected onto an Agilent 1290/SCIEX QTRAP 6500 LC MS/MS system equipped with a C18 reversed phase UHPLC column. The mass spectrometer was operated in negative mode using ESI. The peak area of the individual analyte product ions was measured against the peak area of the product ions of the corresponding internal standards. Quantitation was performed using a weighted linear least squares regression analysis generated from fortified calibration standards prepared concurrently with study samples. LC-MS/MS raw data was collected using SCIEX software Analyst 1.7.3 and processed using SCIEX OS-MQ software v3.1.6. Data reduction is performed using Microsoft Excel for Microsoft 365 MSO.

### Data analysis

For untargeted metabolome data, multiple linear regression, adjusted for sex, age, body weight and study site, was performed. Linear model P values were adjusted for multiple testing using Benjamini-Hochberg method (FDR) [72]. A metabolite was consider significant if FDR <0.10. For SCFAs, ANOVA and Fisher’s LSD multiple comparisons were used to compare group means with alpha of 0.05. To test the null hypothesis that the 2 categorical variables were independent, Fisher’s exact test was used instead. A heatmap was generated where each colored cell on the map corresponded to a concentration value in the data table. Hierarchical clustering was performed using Ward’s method with Euclidean distance. All data analyses and visualizations were performed in the statistical software R (version 4.5.1), GraphPad Prism (version 10.6.1), and MetaboAnalyst (version 6.0).

## Abbreviations

CKD: chronic kidney disease;
2PY: N1-methyl-2-pyridone-5-carboxamide;
SCFA: short-chain fatty acid;
BCFA: branched chain fatty acid;
IRIS: International Renal Interest Society;
NAM: nicotinamide;
PCS: *p*-cresol sulfate;
IS: indoxyl sulfate.

## Declarations

### Ethics approval and consent to participate

The study protocol was reviewed and approved by the Institutional Animal Care and Use Committees (IACUCs) of Nestlé Purina PetCare Company, Ohio State University, Kansas State University, University of Pennsylvania, CanCog Inc. prior to initiation. Informed owner consent was obtained for client-owned cats.

### Consent for publication

All authors reviewed and approved the manuscript and consent to publish.

### Availability of supporting data

All data are available in the Supplementary data files.

### Authors’ contributions

TS performed data analysis and interpretation, wrote the manuscript. JMQ enrolled the cats, collected the fecal samples, and interpreted the results. WHW and LA enrolled the cats and collected the fecal samples. JSS conceived the study and interpreted the results. QL conceived the study, performed statistical analysis, data interpretation and visualization, wrote the manuscript.

## Acknowledgments

The authors would like thank Michael Currier and Heather Brown for laboratory assistance, Dr. Brain Zanghi for assistance with sample collection.

