## Supplemental files for "Fecal untargeted metabolomic and short-chain fatty acid analyses in cats with chronic kidney disease"

Supplementary data

**Table S1** Permutational Multivariate Analysis of Variance (PERMANOVA) on site effects.

|  | F.Model | R2 | pval | p.adj |
| --- | --- | --- | --- | --- |
| CAN vs KSU | 3.7417 | 0.13488 | 0.037 | 0.156 |
| CAN vs OSU | 3.2548 | 0.087366 | 0.05 | 0.156 |
| CAN vs PCC | 3.7593 | 0.066233 | 0.03 | 0.156 |
| CAN vs Penn | 1.1468 | 0.054229 | 0.318 | 0.53 |
| CAN vs Tufts | 1.9706 | 0.11612 | 0.133 | 0.285 |
| KSU vs OSU | 0.25954 | 0.007158 | 0.711 | 0.82038 |
| KSU vs PCC | 3.1484 | 0.054144 | 0.052 | 0.156 |
| KSU vs Penn | 0.75776 | 0.033297 | 0.421 | 0.6315 |
| KSU vs Tufts | 0.059217 | 0.003471 | 0.935 | 0.935 |
| OSU vs PCC | 3.4901 | 0.050958 | 0.032 | 0.156 |
| OSU vs Penn | 0.45508 | 0.014022 | 0.562 | 0.76625 |
| OSU vs Tufts | 0.19541 | 0.007186 | 0.793 | 0.84964 |
| PCC vs Penn | 2.74 | 0.050986 | 0.079 | 0.1975 |
| PCC vs Tufts | 1.6322 | 0.034266 | 0.19 | 0.35625 |
| Penn vs Tufts | 0.37134 | 0.027771 | 0.613 | 0.76625 |

**Table S2.** Significant fecal metabolites with known identifies

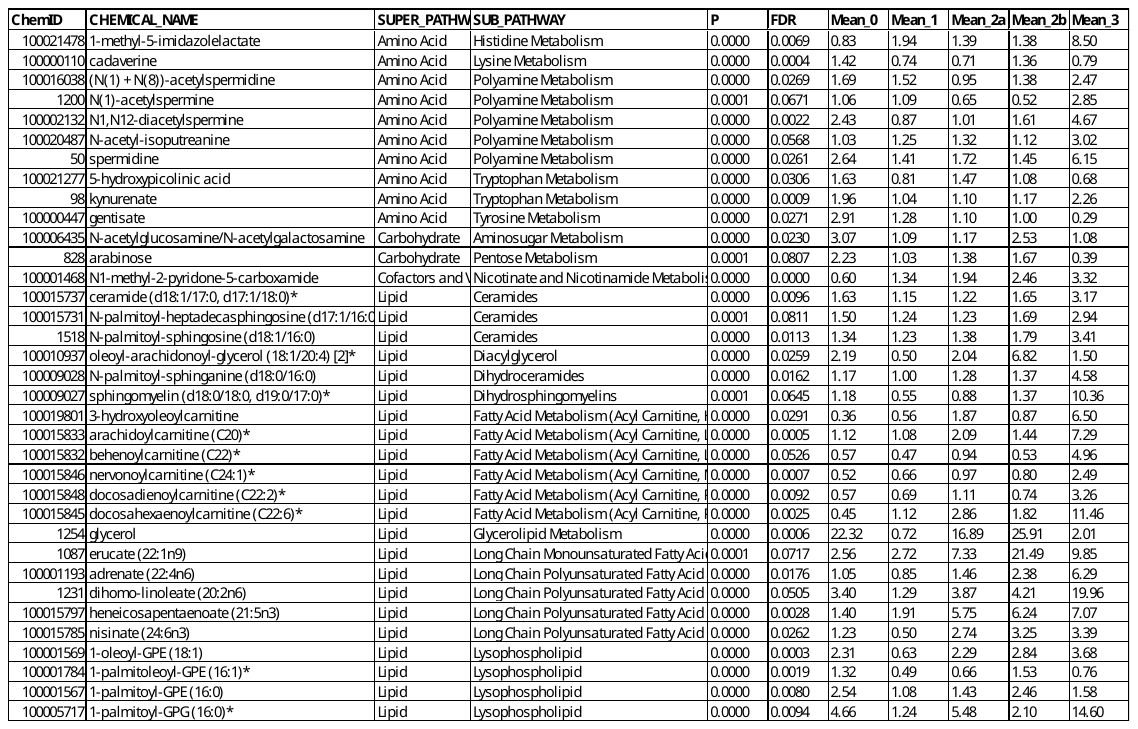

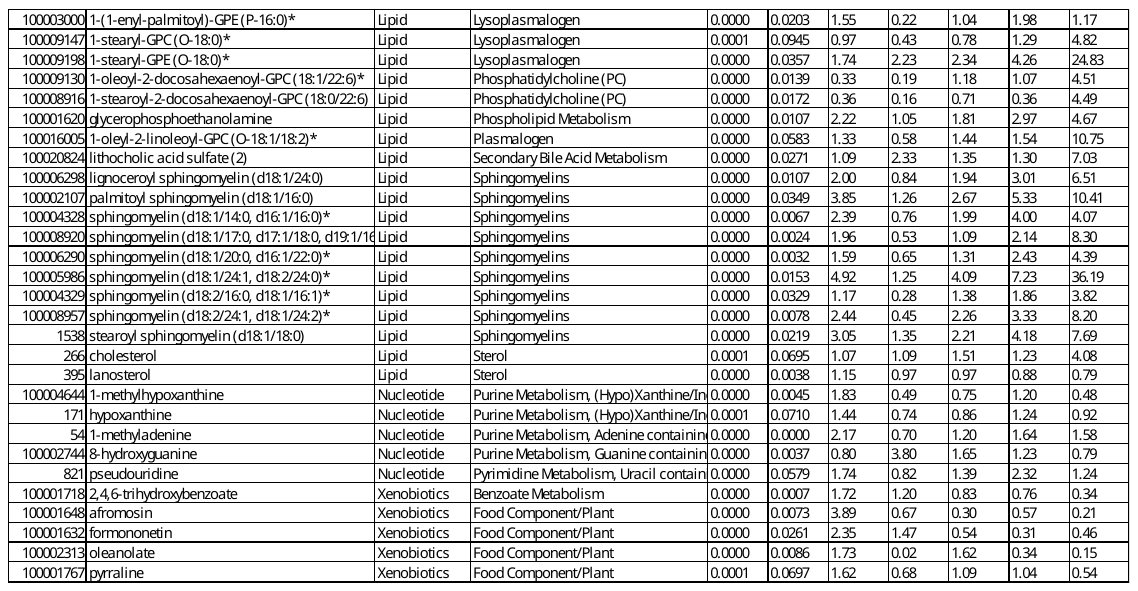

P values were obtained from the multiple linear regression model. FDR were adjusted p values by the Benjamini-Hochberg method.

**Table S3.** Targeted fecal short-chain fatty acids in cats

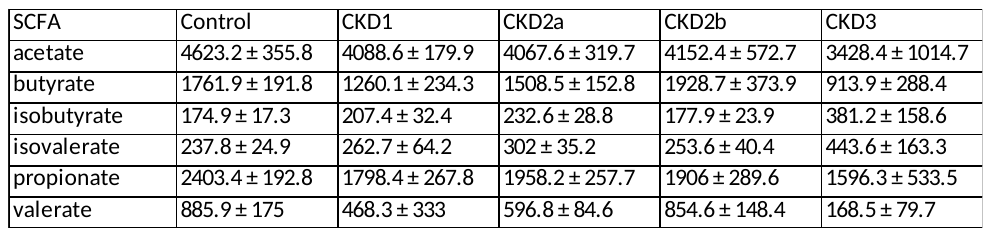

Concentrations were expressed as mean ± SEM in microgram per gram of dry fecal mass (μg/g).

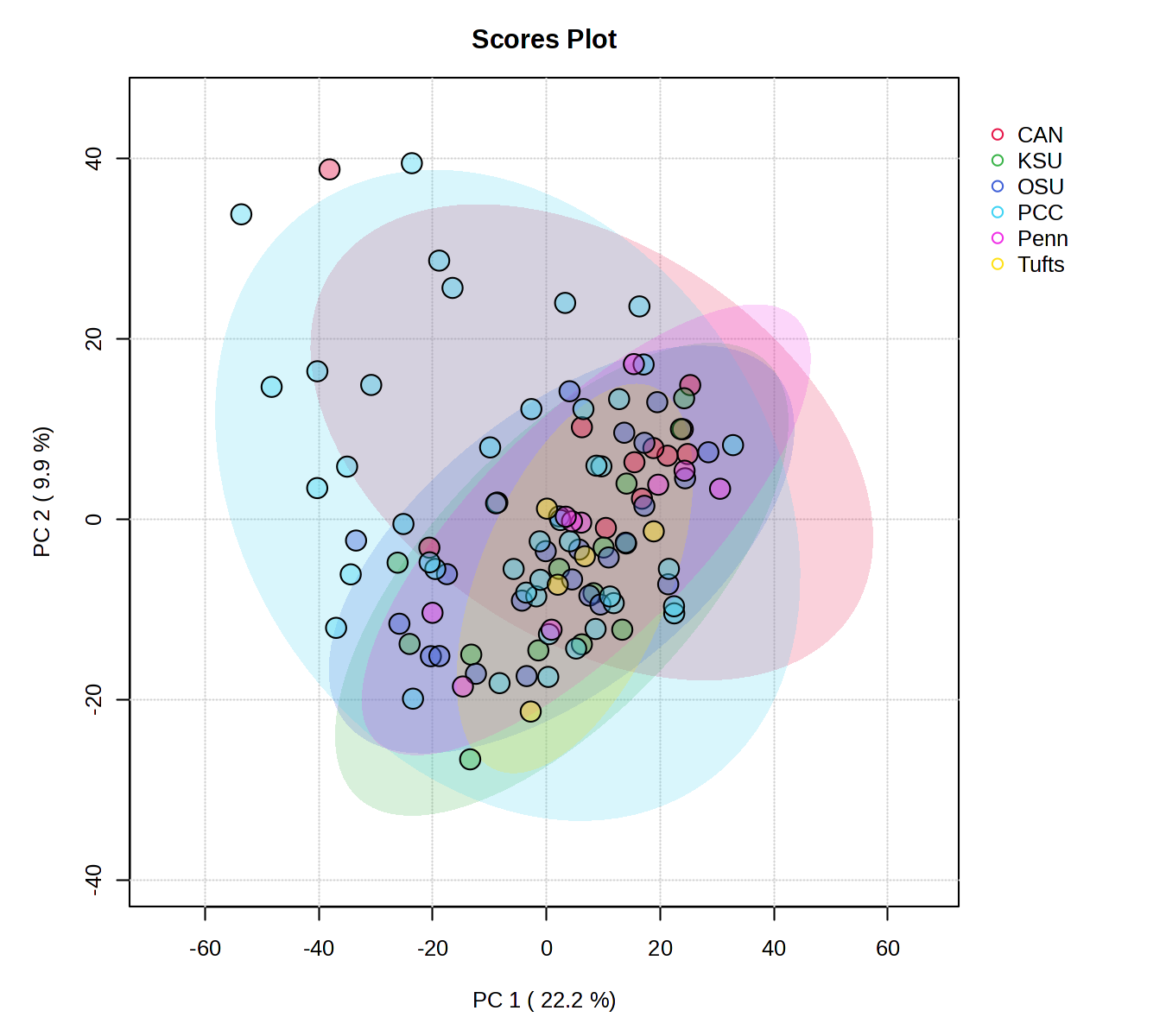

**Fig. S1** Principal Component Analysis of site effects. Pairwise comparisons using Permutational Multivariate Analysis of Variance (PERMANOVA) showed no difference between sites. Also see **Table S3**.

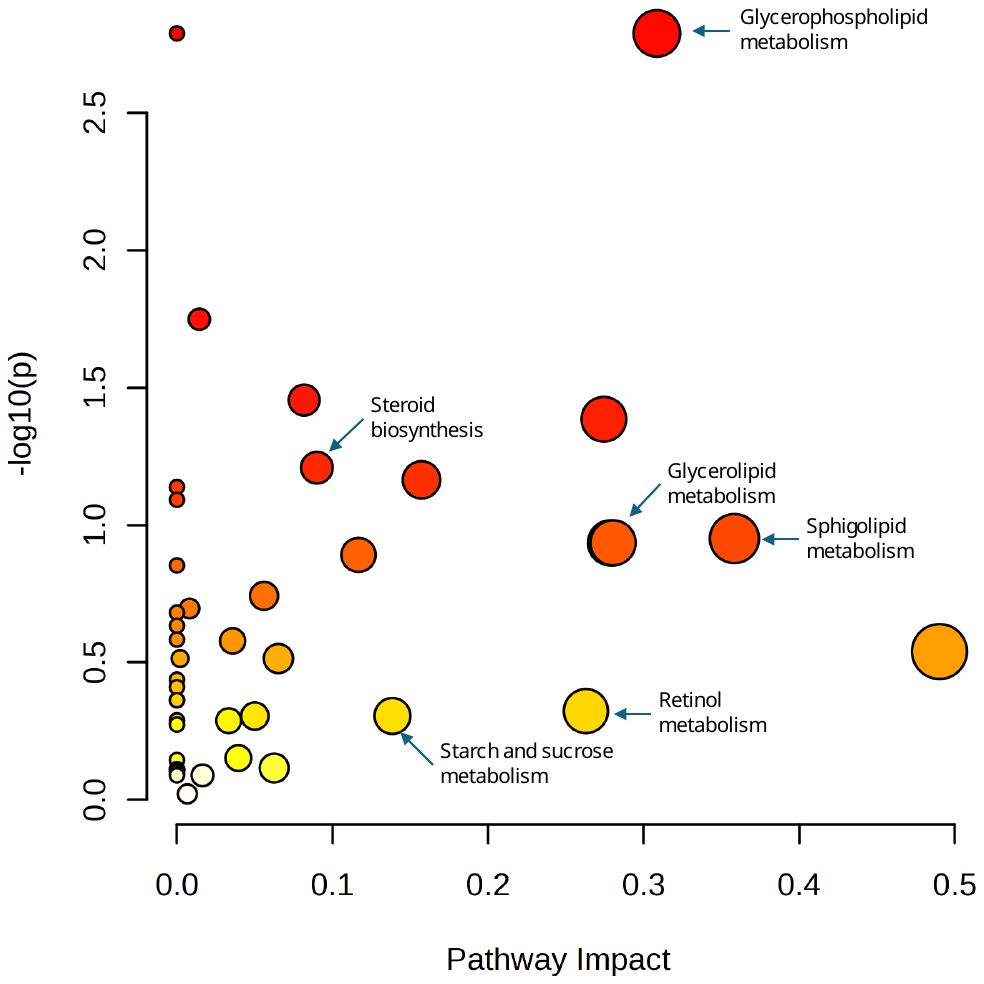

**Fig. S2** KEGG Pathway analysis.
